# Fluctuations in tissue growth portray homeostasis as a critical state and long-time non-Markovian cell proliferation as Markovian

**DOI:** 10.1101/2023.06.07.544008

**Authors:** Natalia G. Lavalle, Osvaldo Chara, Tomás S. Grigera

## Abstract

Tissue growth is an emerging phenomenon that results from the cell-level interplay between proliferation and apoptosis, which is crucial during embryonic development, tissue regeneration, as well as in pathological conditions such as cancer. In this theoretical article, we address the problem of stochasticity in tissue growth by first considering a minimal Markovian model of tissue size, quantified as the number of cells in a simulated tissue, subjected to both proliferation and apoptosis. We find two dynamic phases, growth and decay, separated by a critical state representing a homeostatic tissue. Since the main limitation of the Markovian model is its neglect of the cell cycle, we incorporated a refractory period that temporarily prevents proliferation immediately following cell division, as a minimal proxy for the cell cycle, and studied the model in the growth phase. Importantly, we obtained from this last model an effective Markovian rate, which accurately describes general trends of tissue size. This study shows that the dynamics of tissue growth can be theoretically conceptualized as a Markovian process where homeostasis is a critical state flanked by decay and growth phases. Notably, in the growing non-Markovian model, a Markovian-like growth process emerges at large time scales.

## 1 Introduction

During embryonic development, tissues grow while acquiring patterning, following tightly regulated mechanisms selected after millions of years of evolution [1, 2]. Similarly, regenerating tissues after amputation or ablation expand in an initially explosive and yet controlled growth in those species capable of this trait, such as hydra [3], axolotl [4], planaria [5, 6], zebrafish [7] among many others. In cancer tissues, the explosive tissue growth distinctively progresses lacking some of the controlled mechanisms [8]. The opposite is observed when tissues reduce their size during atrophy, as observed during Sarcopenia [9]. In between these two extreme tissue dynamics, mature tissues reach a final size during adulthood [1]. Hence, like other materials and dynamical systems, the tissues from different species and stages, in physiological and pathological conditions, can be conceptualized in terms of two dynamical phases, growth and decay, separated by an intermediate dynamical state, where tissue size is homeostatically conserved. What determines how tissues transition from one dynamical phase to another has not been yet fully understood.

Tissue growth and decay emerge from the interplay between two cell-based mechanisms: cell death, typically by apoptosis, and cell proliferation. During mammalian embryonic and fetal development, apoptosis transiently predominates over cell proliferation, playing key roles in a broad range of tissues [10]. A similar interplay between both cellular mechanisms was observed in the context of regeneration, where an apoptotic wave triggered by head amputation is crucial for the compensatory proliferation that leads to complete head regeneration in Hydra [11]. This interplay is also present in adult tissues in mammals, an example being the mammalian olfactory epithelium, a self-renewing tissue which maintains its size by continuously replacing apoptotic cells with newly proliferating cells [12]. Disrupture of the balance between cell proliferation and apoptosis is at the heart of all tumor growth where either deregulated proliferation or inhibition of apoptosis dominates [13]. Thus, the transition from the growth to the decay dynamical phases reflects but a transition from a phase dominated by cell proliferation to one in which apoptosis predominates, separated by a state of homeostatic balance between both cellular processes (Fig. 1).

**Figure 1.**
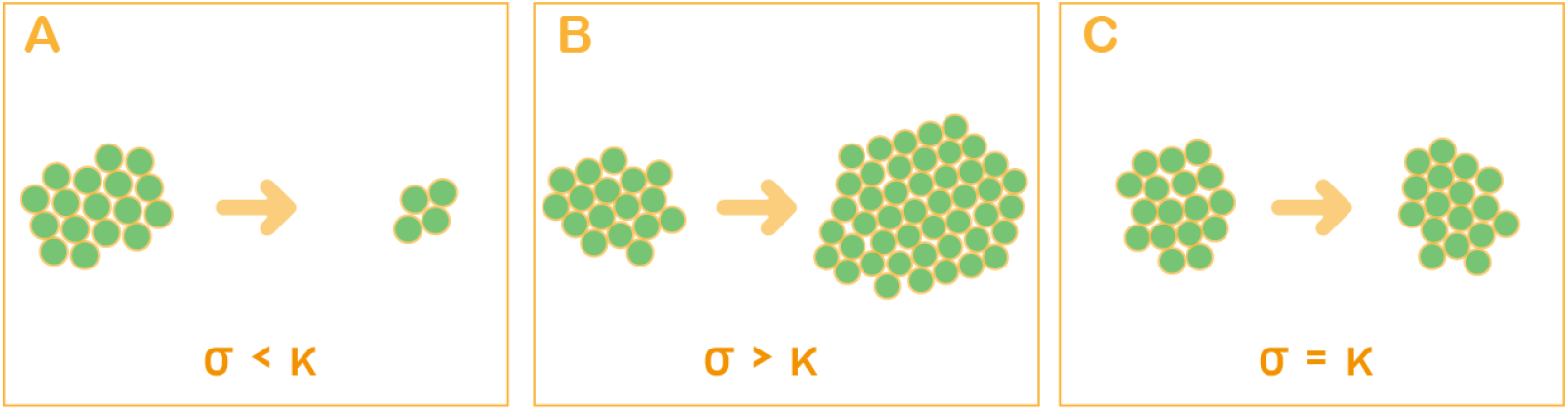
Decay (A) and growth (B) dynamical phases separated by a homeostatic state (C). Schematic illustration of the time evolution of a tissue. (A) Decline phase: the number of cells off the tissue decreases with time. (B) Growth phase: the number of cells increases with time. (C) Homeostatic phase: the number of cells remains approximately constant over time. The arrow represents the time progression. *σ* is the proliferation rate and *κ* is the rate of apoptosis (see Section 2(a)).

Different theoretical approaches have been advanced over the years to model apoptosis, where the emphasis was made on different aspects of the signaling network controlling cell death [14–16]. Cell cycle dynamics was modeled by using stochastic frameworks [17–20]. Based on the seminal stochastic cell cycle model of Smith and Martin[21], a number of studies created very accurate representations of the cell cycle using hypoexponential distributions (see for instance [22, 23]). Accurate mathematical models of the cell cycle were instrumental to understand the impact of cell cycle dynamics on gene expression [24, 25] and tissue development and regeneration [26]. Temporal and sample-to-sample stochastic fluctuations in the number of cells are inherent features of these models. However how these fluctuations impact tissue dynamics during the three dynamical phases and their transitions is not clear.

In this study, we investigated the role of fluctuations in the number of cells on the above mentioned dynamical phases, as well as their transitions, by using an unconstrained Markovian model including stochastic cell proliferation and apoptosis. We analytically calculated the mean, variance and time correlations of the number of cells, which we numerically validate with a computational model. These results allowed us to understand the homeostatic intermediate state as a critical state separating a growth and a decay phases. Then, we adopted a refractory period as a first proxy for the lower bound of the cell cycle length, and we studied the resulting non-Markovian model in the growth phase. Analytical calculations of mean and variance, together with numerical evaluation of time correlations of this extremely synchronized non-Markovian model portray a long-term Markovian dynamics modulated by transient oscillatory features due to the non-Markovian nature of the cell cycle.

## 2 Results

### 2.1 A Markovian model of tissue dynamics displays two dynamical phases separated by a critical state

To investigate the role of fluctuations in the number of cells in the dynamical regimes of growth, decay and homeostasis (Fig. 1), we start with the simplest stochastic model one could think of. We assume a Markovian birth and death model in continuous time, devoid of spatial structure: each cell can undergo proliferation or apoptosis with rates (transition probability per unit time) *σ* and *κ*, respectively. Of note, although it is clear that *σ* and *κ* rates could also represent influx and efflux of migratory cells, cell extrusion, cell differentiation, among other cellular mechanisms, in this study we will arbitrarily focus in cell proliferation and apoptosis. There is no consideration of a cell’s internal state: a cell can proliferate again immediately after a division, always with the same probability rate, and there is no bound on how large the population can grow. The model is a continuous time branching process, and it is completely defined by the so-called master equations [27], a set of coupled differential equations which describe the evolution of the probabilities *P*_*n*_(*t*) that exactly *n* cells are present in the system at time *t*:

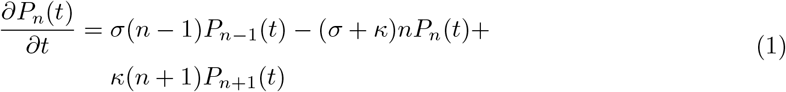

This infinite set of equations can be solved exactly by the method of the generating function (see SM, [28, §2.3.2], and [29, Ch. 1]). From the probabilities *P*_*n*_(*t*) one can compute the mean value of the number of cells ⟨ *n*(*t*) ⟩ and the variance as a function of time, as well as the extinction probability, i.e. the probability that the population vanishes. The state with no individuals is an absorbing state, meaning that if the system reaches this state, it can never leave it. We find (see SM for details)

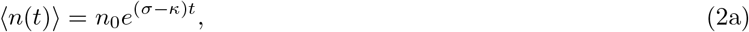

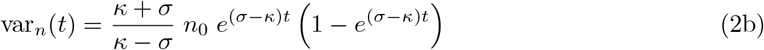

for σ ≠ κ, and

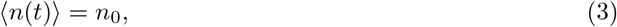

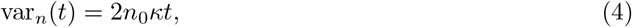

for *σ* = *κ*.

The extinction probability *P*_*e*_, i.e. the probability that the population will reach zero at some point in time, can be computed from the *t* → ∞ limit of *P*_0_(*t*) (see SM):

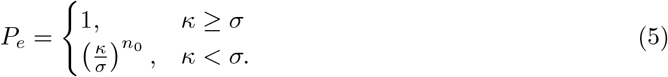

We can thus distinguish two different dynamical phases according to the relationship between the rates of proliferation and apoptosis [28]. If the rate of proliferation is less than that of apoptosis (*σ < κ*), the system is in a decline phase where the average number of cells ⟨ *n*(*t*) ⟩ decreases exponentially (Fig. 2A), while for *σ > κ* we have a growth phase where ⟨ *n*(*t*) ⟩ increases exponentially (Fig. 2C). However, the extinction probability is still finite (Supplementary Fig. 1B) even in the growth phase: a finite fraction of systems will go extinct by chance fluctuations even if the proliferation rate is larger than apoptosis. This is reflected in an exponentially growing variance, as discussed next. Finally, for *σ* = *κ* we have a critical state where the extinction probability is singular and ⟨ *n*(*t*) ⟩ is constant. This critical case can be interpreted as a homeostatic tissue (Fig. 2E), but we stress that unless both *σ* and *κ* are equal to 0, this homeostatic case will have nontrivial dynamics, as can be deduced from the time correlation functions we discuss below.

**Figure 2.**
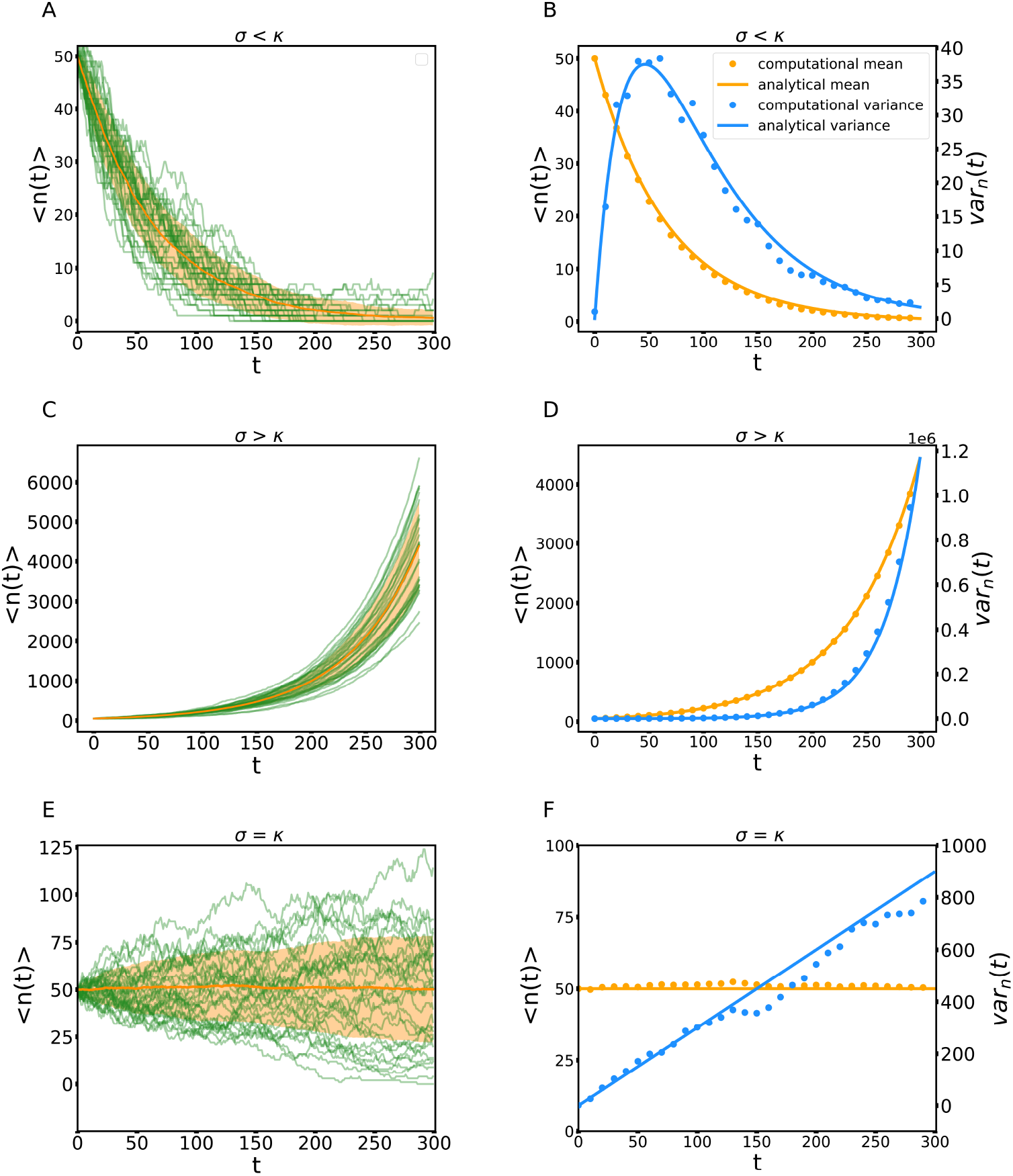
The homeostatic phase operates as a critical system. Time evolution of the mean of the number of cells for 200 samples of a tissue with an initial population of 50 cells, where each cell is assigned a probability of apoptosis (*κ*) and division (*σ*). (A–B) Decline phase. (C–D) Growth phase. (E–F) Homeostatic phase. For panels A, C and E, the green curves are a set of 30 randomly-chosen individual samples, the orange curve is the average of all samples, and the orange shadow gives one standard deviation from the average. For panels B, D and F, in orange the mean and in blue the variance; solid lines correspond to the analytical expressions for the continuous time model, and the points were obtained from numerical simulations of the computational model using 300 samples. (A–B) *σ* = 0.015 and *κ* = 0.03, (C–D) *σ* = 0.03 and *κ* = 0.015, (E–F) *σ* = *κ* = 0.03. for *σ* ≠ *κ* (we turn to the critical *σ* = *κ* case below). This expression justifies the intuitive feeling that the different samples are simply exponentials that amplify the fluctuations that occur when the population is still small: the product *n*(*t*_1_)*n*(*t*_2_) is on average *n*^2^(*t*_1_) multiplied by an exponential with the same rate as ⟨ *n*(*t*) ⟩. This effect is also found in other models with exponential growth, such as epidemic spreading in its early stages [31].

To validate our analytical results, we compared the analytical expressions with numerical results of a computational discrete-time version of the models (individual curves in Fig. 2 A,C,E) and obtained a good agreement (Fig. 2 B,D,F). In the decline phase (Fig. 2A and B) the mean decreases exponentially, while the variance initially increases, but after reaching a maximum it goes to zero exponentially. Inspection of the individual stochastic samples shows that several of them reach the absorbing state with no cells, a manifestation of the fact that the extinction probability is 1. In the growth phase *σ > κ* (Fig. 2 C and D) both the mean and variance are exponentially increasing, Eqs. (2). The extinction probability is very low (9·10^*−*16^) for the values of the parameters chosen in the simulation, and accordingly none of our samples go extinct. The variance grows rapidly, but this reflects inter-sample fluctuations, rather than temporal fluctuations within a single sample: the growth of a single sample exhibits rather small fluctuations around an exponential curve (Fig. 2C). The fluctuations around the mean are mainly due to the fluctuations that take place at short times. One can say that once growth “kicks off”, then it proceeds exponentially, and the different samples just look like time-shifted versions of one-another (Supplementary Fig. 2). The growing variance does not reflect the fact that it is extremely unlikely that two growth curves will meet again once they have diverged. We will come back to this point when discussing the time correlation function. Finally, the homeostatic case (*σ* = *κ*) shows large fluctuations in the individual samples, even though the average number of cells is constant in time (Fig. 2E). The variance grows linearly (Fig. 2F), reflecting fluctuations among samples. Unlike the growth phase, the population curves of individual samples show complicated shapes, with periods of growth and decrease, and do not look lie time-shifted copies. The extinction probability is 1, but while many samples go extinct relatively fast, there are a few that grow significantly before going to the absorbing state. This case can be described as critical, in the sense that it is the boundary between the decline and growth phases, and that the extinction probability as a function of *σ/κ* is singular at *σ/κ* = 1, in the sense that it is not analytic in any neighborhood of *σ/κ* = 1 because it equal to the constant 1 for *σ/κ <* 1, and non-constant for *σ/κ >* 1. It also shares with critical systems the fact that the correlation time is infinite (more on that below). Finally, it is important to remark that the homeostatic state is not stationary, because although the mean number of cells is time-independent, the variance is growing with time. In the terminology of stochastic processes [27], it is only stationary up to first order.

### 2.2 Time correlation of the Markovian model confirms that the homeostatic case is a critical state

A deeper understanding of the different dynamical states can be obtained from the time correlation function of number of cells,

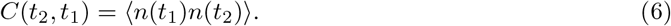

For this model this function can be obtained analytically as (see SM and refs. [29, 30]):

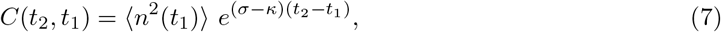

To measure how fluctuations decorrelate, it is more useful to compute the auto-covariance, or connected correlation function as it is known in the physics literature,

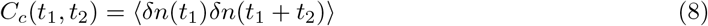

because unlike (6) this will vanish when *n*(*t*_1_) and *n*(*t*_2_) are uncorrelated. This is readily obtained as which shows that correlations decay exponentially for *σ < κ* (Fig. 3A), but fluctuations never fully decorrelate if *σ > κ* (Fig. 3C). Indeed, in the last case the connected correlation is actually monotonically increasing for increasing *t*_2_ − *t*_1_ (and diverges for *t*_2_ − *t*_1_ → ∞). This growth with time might give the impression that correlation is stronger for larger *t*_2_ −*t*_1_, contrary to what one expects. However, this growth is actually reflecting the fact that the *amplitude* of the fluctuations is growing (recall the variance diverges exponentially). To get rid of this effect we can define a normalized correlation function

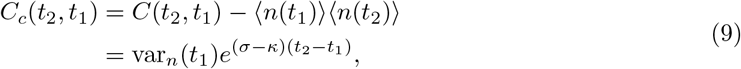

which is nothing but the Pearson correlation coefficient of *n*(*t*_1_) and *n*(*t*_2_), and is guaranteed to lie in the interval [−1, 1] (with 1 holding for perfect linear correlation). It is equivalent to computing the connect<u>ed cor</u>relation of the time series normalized with the instantaneous standard deviation, i.e. 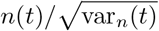. This quantity is monotonically decreasing in both dynamical phases (Fig. 3B, 3D). However, for *σ > κ* it never vanishes, but tends, for *t*_2_ → ∞ at fixed *t*_1_, to

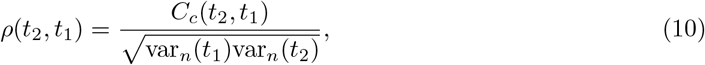

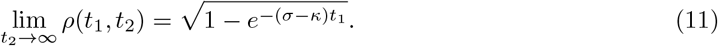

**Figure 3.**
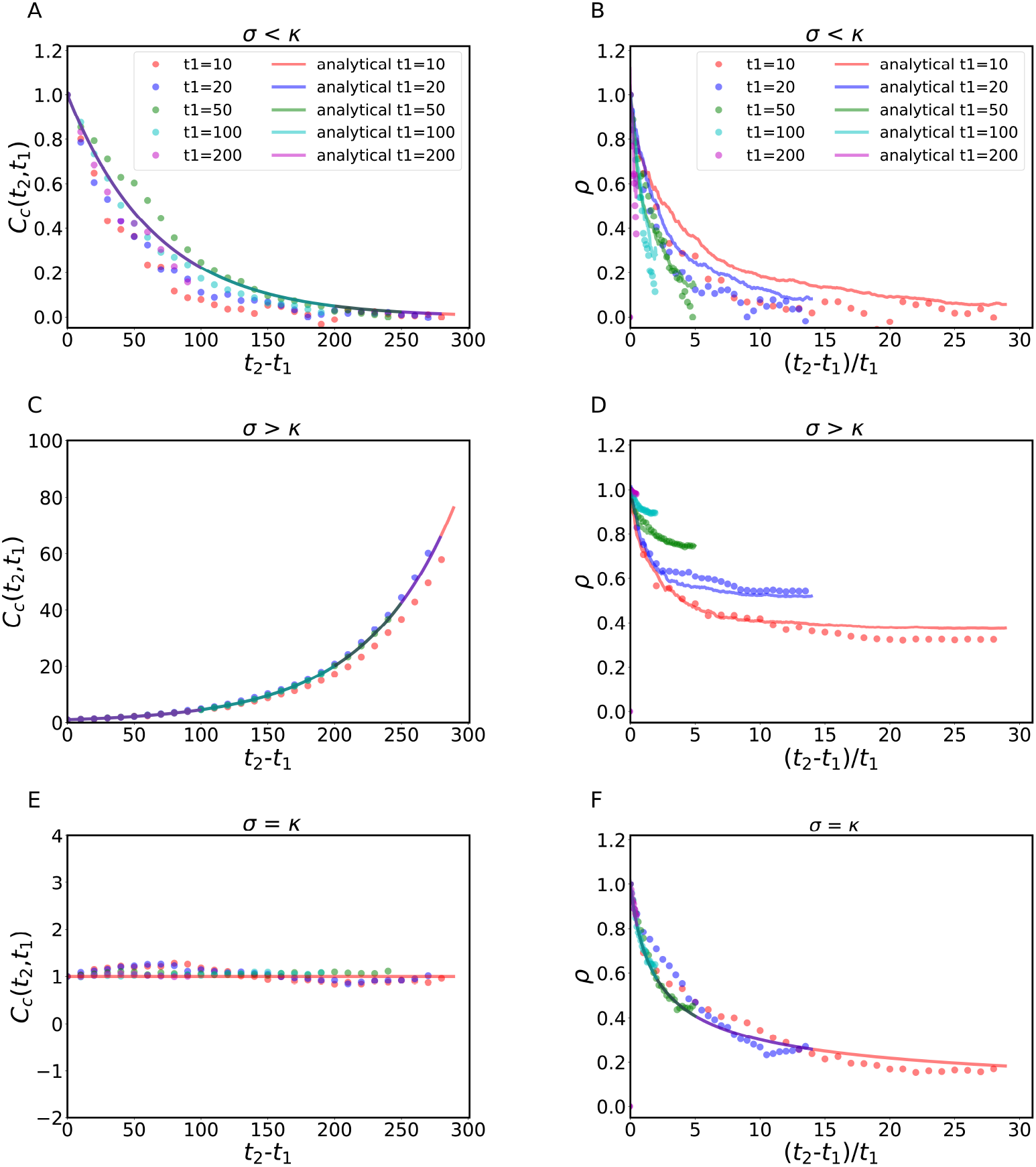
Fluctuations of the Markovian model decorrelate in time in the three dynamical regimes. There is quantitative agreement between analytical expressions (solid lines) and numerical calculations using the computational model (dots) of the connected (A, C, E) and normalized (B, D, F) time correlations between times *t*_1_ and *t*_2_ for the decline (A, B), growth (C, D), and homeostatic cases (E, F). In the decay phase (A, B) fluctuations decorrelate. In the growth phase, we observe that the connected correlation grows (C), due to an increase of the amplitude of the fluctuations, and the normalized correlation (D) shows that fluctuations decorrelate, but not fully, and the limit *t*_2_→∞ is finite (higher the higher *t*_1_). This is the rigorous statement of the intuitive fact that when growth “kics off” in a sample, it just keeps growing exponentially. Finally, the homeostatic case displays a constant connected correlation (E) and a power-law decay of the normalized correlation, with critical exponent 1*/*2 (and consequently an infinite correlation time).

That is, for large *t*_1_, fluctuations stay strongly correlated throughout. In other words, if at a given time, the population of one sample is larger than that of another sample, it is highly likely that the first sample will continue to be larger than the second sample in the future (while both samples grow exponentially). A final note for these noncritical cases is that, for *σ < κ*, the correlations decay exponentially and one can define a correlation time

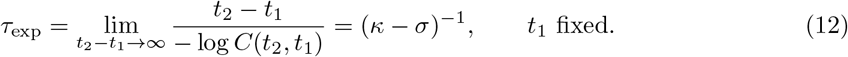

Turning now to the critical case, the connected correlation is

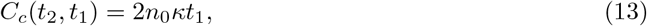

which is a constant for a fixed *t*_1_ (Fig. 3E). This result underlines again that the system is not stationary, even though ⟨ *n*(*t*)⟩ is time-independent. The fact that the correlation function is constant for fixed *t*_1_ can be understood by noting that in this case *n*(*t*) behaves roughly like a random walker (with an absorbing wall at *n* = 0), because at each time step it is equally likely that *n* will increase or decrease. This is consistent with the facts that the variance is linear in time, and that the correlation function depends only on the shortest time [32]. Of note, the analogy with the random walk is not a perfect mapping, however, since the amplitude of the random variation of *n* at each time step is proportional to *n* itself.

The normalized (Pearson) correlation does decay, but very slowly, as a power law (Fig. 3F),

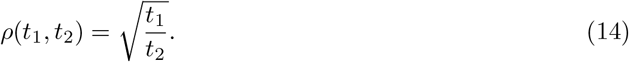

From this result, we see that the relaxation time at fixed *t*_1_ is infinite, another feature of critical systems [28].

### 2.3 Long-term mean and variance of the non-Markovian model converge to those of the Markovian model

The simplicity of the Markovian model presented in previous sections allowed us to analytically derive the expressions for mean, variance, connected correlation and Pearson correlation coefficients of the number of cells. Nevertheless, from the biological perspective this model is clearly oversimplified and a crucial aspect in particular is disregarded: the existence of a cellular cycle. The Markovian model assumes that a cell is always equally likely to divide or die, independently of its position along the cell cycle or age. This is of course unrealistic, as a cell will not divide unless it sequentially goes through the G1, S, G2 phases and reaches the end of M phase. We thus decided to consider a model, also in continuous time, that incorporates such an effect. Disregarding apoptosis for simplicity, we consider a proliferation rate

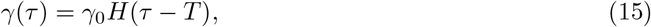

where H is the Heaviside step function and *τ* is the cell’s age, i.e. the time elapsed since the last cell division. In other words, cell division is forbidden when the age of the cell is less than *T*, otherwise the proliferation rate is constant as in the previous model. Of note, the proliferation rate can be considered an effective rate, featuring the balance between proliferation and apoptosis. For the sake of our argument, we will arbitrarily assume that the apoptosis rate in our system is zero. Hence, *T* can be considered as a lower bound of the cell cycle length. This choice makes the model non-Markovian, because the proliferation probability (transition probability in the language of stochastic processes) does not depend only on the number of cells, but also on an internal and continuous state variable (the position along the cell cycle or age). Non-Markovian processes are much harder to solve than Markovian ones, but in this case we can use the method described in reference [27] (details in SM) to obtain the mean and variance of the population. The mean population is found as (see SM)

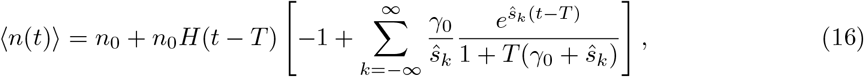

where *ŝ*_*k*_ are the poles of the Laplace transform of ⟨ *n*(*t*)⟩, given by

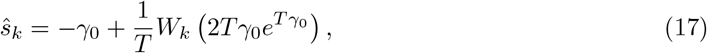

with *W*_*k*_(*x*) the *k*-th branch of Lambert’s function (also known as product log). *ŝ*_0_ is real, but the rest of the poles appear in complex-conjugate pairs, with 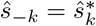.

This analytical expression is in agreement with the mean computed from the numerically simulated discrete-time version of the model (Fig. 4A). More importantly, the general qualitative features (long-term exponential growth, large sample-to-sample variations, high correlations within a sample) are the same as in the Markovian case. The main difference is the appearance, at times not too large, of some oscillations around the exponential curve (Fig. 4A). These oscillations are due to the fact that, when the growth rate constant is larger than the inverse of the refractory period *T*, there are surges of proliferation at times that are multiples of *T*. To see this, consider the case γ ≫1*/T* : at *t* = 0, cells are placed at the beginning of the refractory period, and thus they cannot proliferate for *t < T*. From *t* = *T*, proliferation starts, but since γ is very large, most cells reproduce almost right away, and there is a step-like increase of the average number of cells to twice the initial number. Hence most of the cells are refractory until about *t* = 2*T*, when there is a second step-like increase and so on. However, proliferation is not quite simultaneous, so that as time grows the refractory periods de-synchronise and oscillations tend to disappear. This can be seen analytically from the solution, which asymptotically behaves like

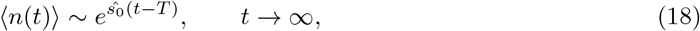

because *ŝ*_0_ is the pole with the largest real part. Thus at large times oscillations vanish and one recovers a pure exponential growth. The last expression also shows that the effective exponential rate of growth is *ŝ*_0_. We have also computed the variance as a function of time. The analytical expression is more complicated, involving a triple infinite sum (full expression in SM). The expression is rather tricky to evaluate numerically due to cancellations among different terms of the series. The analytical expression was fully recapitulated by the numerical simulations (Fig. 4B). The above-mentioned effect, namely the tendency of proliferation to proceed in “waves” due to presence of a refractory period, also causes oscillations in the time derivative of the variance and, for large enough γ, in the variance itself. More precisely, when γ ≫1*/T*, the variance oscillates quite strongly at short times, nearly vanishing at times that are approximately multiples of the refractory period *T* (Fig. 4B). This is due to the fact that since γ is large, most cells proliferate very soon after the refractory period elapses. This is also clear from the curves showing the dispersion of the average number of cells among different simulation samples (Supplementary Fig. 3). The period 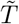 of the oscillations can be obtained approximately from the imaginary part of *ŝ*_1_, which is the oscillation that dominates at intermediate times,

**Figure 4.**
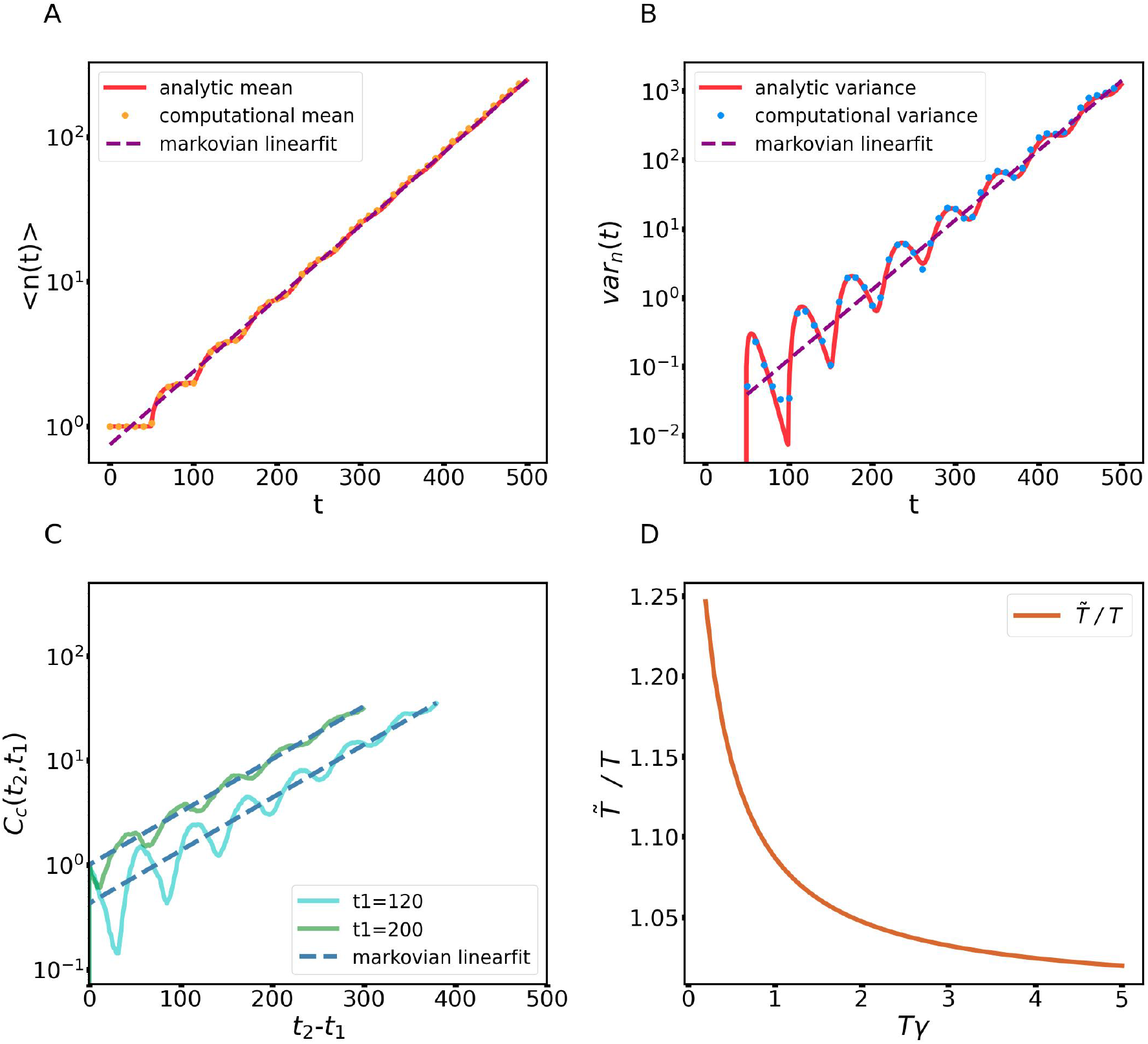
The Markovian model is a good approximation of the non-Markovian model in the growth phase at long times. A) The mean of the cell number for the non-Markovian model analytically obtained (red solid line) agrees with the numerical calculation (orange dots). The linear fit to the logarithm of the cell number mean (violet dashed line) yields a slope which coincides with the first non-trivial pole of the Laplace transform of ⟨ *n*(*t*) ⟩ (*ŝ*_0_ in Eq. 17). B) The analytical variance (red solid line) agrees with the variance computed over 200 samples of the computational model (blue dots). The linear fit to the logarithm of the variance (violet dashed line) gives a slope that coincides with 2*ŝ*_0_. C) The connected time correlation function between times *t*_1_ and *t*_2_ vs. *t*_2_ −*t*_1_ shows oscillations superimposed on an exponential curve with rate *ŝ*_0_. Solid lines are *C*_*c*_(*t*_1_, *t*_2_) for two values of *t*_1_ and dashed lines are linear fit with slopes fixed to *ŝ*_0_. D) Approximate period of oscillations 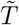 (as measured by the imaginary part of the first complex pole *ŝ*_1_), in units of *T*. When the proliferation rate is much larger than 1*/T*, the period 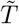 is essentially equal to *T*.

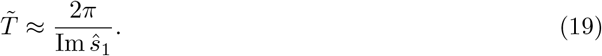

It can be checked that in effect 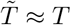 when γ ≫ 1*/T* (Fig. 4D).

Finally, we have computed the time correlation functions for this case. This we have only done numerically, for the discrete-time computational model. The connected correlation function *C*_*c*_ and the normalized (Pearson) correlation share the most important qualitative features with those of the Markovian case: *C*_*c*_*c* grows exponentially with time (Fig. 4C), while *ρ* decays slowly, approaching a finite value at large times. The interpretation of these features is the same as before, namely that the system stays correlated asymptotically because if a chance fluctuation carries the population of some sample above that of another sample of another time, the fast (exponential) growth makes it very likely that the situation will stay the same for all future times. The new feature of the non-Markovian case is the presence of (damped) oscillations superimposed over the mentioned trends. The origin of these oscillations can again be traced back to the refractory period *T*.

## 3 Discussion

We have presented a detailed study of the dynamics of two minimal growth models, a Markovian and a non-Markovian model, the second of which incorporates the cell cycle through a refractory period where cell division is forbidden. The Markovian growth model is actually the well-known continuous time binary branching process, closely related to the Galton-Watson problem [27–30, 33]. The homeostatic case has also been discussed in ref. [34] in the context of epidermal tissue dynamics. In the Markovian model, time correlations in the growth phase and in the homeostatic case (the point in parameter space separating growth from decay) are particularly interesting. In the growth phase, fluctuations stay forever correlated, and samples found to have a larger population than other samples at a given point in time will likely stay larger in the future. We have shown that the homeostatic case is critical: time correlations decay but the correlation time is infinite, and the variance grows indefinitely with time, showing that fluctuations are strong in this state despite the mean number of cells being time-independent.

The Markovian model is clearly too simple for a realistic model of cell growth, the lack of cell cycling being its most obvious oversimplification. The cell cycle is hard to treat mathematically, because its inclusion makes models non-Markovian: the division probability has to depend on an internal real variable, namely the time since a cell has last divided. For this reason, cell cycle is often neglected in models of tissue growth, although it can be of importance for an accurate description of key cellular processes, as elegantly illustrated by Perez-Carrasco and colleagues [24]. Indeed, this study explicitly analyzed the error in the mRNA transcription rate estimation owing to neglecting cell cycle variability and replication.

We here developed a non-Markovian model for the growth phase, which incorporates a näive cell cycle in the form of a refractory period: the proliferation rate is *γ*, but proliferation is forbidden if the cell age is less than *T*, a situation is often approximated with multi-stage Markovian models (e.g. [23]). The same model was considered in [34], where it was solved numerically, focusing on clone size distributions. Here we have instead found an exact solution for the mean and variance in this minimal non-Markovian case, and have shown that the new feature is the appearance of oscillations that are superimposed on the same general trends of the Markovian case. Since the oscillations are damped, for large times the two models are mostly qualitatively indistinguishable. Although the analytical expressions for the non-Markovian case are somewhat awkward to handle, knowing the refractory period and the non-Markovian rate it is possible to calculate an effective asymptotic Markovian rate (using expression (17) with *k* = 0). This allows to make use of the simpler Markovian formulae, Eqs. (2) which furnish a rather good approximation for the non-Markovian growth at large times. Apart from the oscillations, these expressions, with the effective Markovian rate, accurately describe the general trends of the mean, variance, and correlation functions. A similar comparison between the Markovian and non-Markovian case was done, for the case of stochastic gene expression, by [25].

We have also found an approximate expression for the period of the oscillations (19). The damping of the oscillation is due to a de-synchronization effect: when starting from a single cell (or a group of synchronized cells), if the reproduction rate is larger than the inverse of the refractory period, proliferation occurs in “waves”, i.e. most cells proliferate shortly after the refractory period, then cannot proliferate during a new refractory period, and so on. However, since proliferation is stochastic, after several divisions, the cell cycles de-synchronize and oscillations die out. This can observed experimentally by using markers monitoring the transit through the cell cycle, like Fluorescent Ubiquitination-based Cell Cycle Indicator (FUCCI, [35]). A clear example showing high synchronization of cells displayed by FUCCI in vitro can be found in the study of Vittadello and colleagues [36]. A more recent example showing how FUCCI reveals cell synchronization in vivo during regeneration of the spinal cord in axolotls is our study of Cura Costa and colleagues [26]. Importantly, in both examples synchronization revealed by FUCCI was recapitulated with a non-Markovian model of the cell cycle ([36], [26]).

Finally, it should be pointed out that although the non-Markovian model improves on the Markovian one by including (crudely) some internals of the cell’s functioning, both models share two important limitations, namely the fact that they ignore interactions among cells, and that they include no spatial information (in particular they completely ignore cell motility). It is known that motility reduction and mechanical interactions due to crowding influence cell cycle progression and have an important role in controlling proliferation [37, 38]. Thus the present models can only be valid for times such that they do not predict large changes in cell density.

## 4 Conclusions

The Markovian model here studied can reproduce the main features of the dynamics of a growing tissue at long times, even though it is a minimal model that does not take into account actual cell cycling. An advantage of this model is that we could derive exact and manageable analytical expressions for the mean, variance, connected connections and Pearson coefficient of the population. However, to study tissue growth at shorter times, one must take into account the cell cycle explicitly, which introduces oscillations in the mean and variance of the population. This is a more realistic approximation, but harder to treat analytically. Nevertheless, We have obtained exact expressions for the mean and variance, though somewhat unwieldy for this model. We have shown that the non-Markovian model behaves qualitatively like the Markovian model for large times due to de-synchronization of cell cycles, and have given an expression for the effective exponential growth rate.

## 5 Methods

We have considered two mathematical models of tissue growth, devoid of spatial structure. The first model is a Markovian birth and death model in continuous time, the second one simulates the cellular cycle by incorporating a refractory period during which the cells cannot divide, which makes the model non-Markovian. For the Markovian model, we have solved the master equations with the generating function technique [27–29] to obtain analytical expressions for the mean, variance and time correlation function of the number of cells, as explained in detail in the Supplementary Materials. In the non-Markovian case we were able to obtain analytical expressions for mean and variance by solving a nonlinear integral equation obeyed by the generating function of this branching process [27], as detailed in the SM.

We contrasted the analytical results with numerical simulations of the same models cast in discrete time. The models were implemented in Python (SM Sec. III). To do the comparison, the transition probabilities *P*_*σ*_, *P*_*κ*_ used in the discrete case must be related to the transition *rates* used in continuous time by

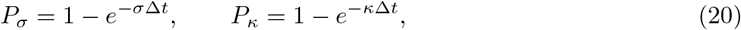

where ∆*t* is the discrete time step. We simulated 200 samples of the stochastic dynamics generated with the computational models and computed the mean and variance of the number of cells, together with the connected and the Pearson correlation.

## 6 Acknowledgements

We thank the members of the Chara and Grigera labs for their invaluable comments on this study.

