## Supplementary Material for "Fluctuations in tissue growth portray homeostasis as a critical state and long-time non-Markovian cell proliferation as Markovian"

### Suplementary Material

Natalia G. Lavalle
*Instituto de Física de Líquidos y Sistemas Biológicos (IFLySiB) — Universidad Nacional de La Plata and CONICET, Calle 59 n. 789, B1900BTE La Plata, Argentina*

Osvaldo Chara†*
*School of Biosciences, University of Nottingham,*
*Sutton Bonington Campus, Nottingham LE12 5RD, UK*
*Instituto de Tecnología, Universidad Argentina de la Empresa, Buenos Aires C1073AAO, Argentina*

Tomás S. Grigera†*
*Instituto de Física de Líquidos y Sistemas Biológicos (IFLySiB) — Universidad Nacional de La Plata and CONICET, Calle 59 n. 789, B1900BTE La Plata, Argentina*
*CCT CONICET La Plata, Consejo Nacional de Investigaciones Científicas y Técnicas, Argentina*
*Departamento de Física, Facultad de Ciencias Exactas, Universidad Nacional de La Plata, Argentina*
*Istituto dei Sistemi Complessi,*
*Consiglio Nazionale delle Ricerche,*
*via dei Taurini 19, 00185 Rome, Italy*

† *These authors contributed equally to this study*

**Authors for correspondence:*
*Osvaldo Chara* **
*Tomás S. Grigera* **

#### CONTENTS

I. Markovian Model 3
A. Mean and variance 4
B. Correlation Function 4

II. Non-Markovian model 5
A. Population mean 6
B. Variance 7
C. Numerical evaluation of mean and variance 7

III. Numerical simulations 7

IV. Supplementary figures 8

References 9

#### I. MARKOVIAN MODEL

The simple Markovian model of tissue growth is a continuous time binary branching process closely related to the Galton-Watson problem. Here the time is continuous and cells can undergo proliferation or apoptosis. The spatial distribution of cells is ignored, and the only dynamical variable is the number of cells $n(t)$. We assume that each individual initiates a family of descendants, that all families are equivalent (no mutations), and that families do not interact (asexual reproduction). The rates of proliferation, $\sigma$, and apoptosis, $\kappa$, for a single cell are constant. In particular they are independent of the age of the cell: this is what makes the model Markovian, and amounts to ignoring the existence of a cell cycle (and of cell growth and ageing). Then transitions from a state with $n$ cells to a state with $n+1$ cells occur with rate $n\sigma$, while transitions $n \to n-1$ have rate $n\kappa$, and the state $n = 0$ is absorbing. Calling $P_n(t)$ the probability of finding $n$ cells at time $t$, its time evolution is given by the set of master equations,

$$\frac{\partial P_n(t)}{\partial t} = \sigma(n-1)P_{n-1}(t) - (\sigma+\kappa)nP_n(t) + \kappa(n+1)P_{n+1}(t), \qquad \text{for } n \geq 2, \tag{I.1a}$$

$$\frac{\partial P_1(t)}{\partial t} = -(\sigma+\kappa)P_1(t) + 2\kappa P_2(t), \tag{I.1b}$$

$$\frac{\partial P_0(t)}{\partial t} = \kappa P_1(t). \tag{I.1c}$$

This set of infinitely many ordinary differential equations can be solved by the method of the generating function [1, 2](the model has also been solved by introducing a suitable field theory [3]). The generating function is defined as

$$g(x,t) = \sum_{n=0}^{\infty} x^n P_n(t), \tag{I.2}$$

from which the $P_n(t)$ can be obtained by differentiation, and in terms of which all moments of $n(t)$ can be written. Here we are interested in the mean and variance,

$$\langle n(t)\rangle = \sum_{n=1}^{\infty} nP_n(t) = \left.\frac{\partial g(x,t)}{\partial x}\right|_{x=1}, \tag{I.3}$$

$$\begin{aligned} \mathrm{var}_n(t) &= \langle n^2(t)\rangle - \langle n(t)\rangle^2 \\ &= \left.\frac{\partial g(x,t)}{\partial x}\right|_{x=1} + \left.\frac{\partial^2 g(x,t)}{\partial x^2}\right|_{x=1} - \left(\left.\frac{\partial g(x,t)}{\partial x}\right|_{x=1}\right)^2. \end{aligned} \tag{I.4}$$

The generating function obeys the equation

$$\begin{aligned} \frac{\partial g(x,t)}{\partial t} &= \sum_{n=0}^{\infty} \left[\sigma x^{n+1} - (\sigma+\kappa)^n + \kappa x^{n-1}\right] nP_n(t) \\ &= \kappa(1-x)\left(1 - \frac{\sigma}{\kappa}x\right)\frac{\partial g(x,t)}{\partial x}, \end{aligned} \tag{I.5}$$

subject to the boundary condition

$$g(1,t) = \sum_{n=0}^{\infty} P_n(t) = 1, \tag{I.6}$$

which can be solved by the method of characteristics [1]. The solution is

$$g(x,t) = \left[\frac{(\kappa-\sigma x)e^{(\kappa-\sigma)t} - \kappa(1-x)}{(\kappa-\sigma x)e^{(\kappa-\sigma)t} - \sigma(1-x)}\right]^{n_0}, \tag{I.7}$$

for $\sigma \neq \kappa$, and

$$g(x,t) = \left[\frac{1+(1-x)(\kappa t-1)}{1+(1-x)\kappa t}\right]^{n_0} \tag{I.8}$$

for $\sigma = \kappa$.

##### A. Mean and variance

From equations (I.3), (I.4), (I.7), and (I.8) one obtains the mean and variance at time $t$ given that there are $n_0$ cells at time $t = 0$ as

$$\langle n(t)\rangle = n_0 e^{(\sigma-\kappa)t}, \tag{I.9}$$

$$\mathrm{var}_n(t) = \frac{\kappa+\sigma}{\kappa-\sigma}\, n_0\, e^{(\sigma-\kappa)t}\,(1-e^{(\sigma-\kappa)t}), \tag{I.10}$$

for $\sigma \neq \kappa$, and

$$\langle n(t)\rangle = n_0, \tag{I.11}$$

$$\mathrm{var}_n(t) = 2n_0\kappa t, \tag{I.12}$$

for $\sigma = \kappa$.

Eqs. (I.7) and (I.8) also allow to compute the extinction probability $P_e$. Since $n(t) = 0$ is an absorbing state, the probability that a sample is extinct at time $t$ is just $P_0(t)$ (but note it may have become extinct at an earlier time). This is given by $g(0,t)$:

$$P_0(t) = \left[\frac{e^{(\kappa-\sigma)t}-1}{e^{(\kappa-\sigma)t}-\frac{\sigma}{\kappa}}\right]^{n_0}, \qquad \text{if } \sigma \neq \kappa \tag{I.13}$$

$$P_0(t) = \left[\frac{\kappa t}{1+\kappa t}\right]^{n_0}, \qquad \text{if } \sigma = \kappa. \tag{I.14}$$

The asymptotic extinction probability, i.e. the probability that a sample becomes extinct at any point in time, is obtained by taking $t \to \infty$, which yields Eq. (2.5) of the main text.

##### B. Correlation Function

The time correlation function is written in terms of the conditional probabilities $P(n_2t_2|n_1t_1)$ of finding $n_2$ cells at time $t_2$ given that there were $n_1$ cells at $t_1$,

$$C(t_2,t_1) = \langle n(t_1)n(t_2)\rangle = \sum_{n_1,n_2=0}^{\infty} n_1 P(n_2t_2|n_1t_1) n_2 P(n_1t_1|n_0t_0), \tag{I.15}$$

and the conditional probabilities are obtained from the master equation with suitable initial conditions [4]. In this case is suffices to observe that the mean at $t_2$, given that there were $n_1$ cells at $t_1$ is

$$\langle n(t_2)\rangle|_{n_1,t_1} = \sum_{n_2=0}^{\infty} n_2 P(n_2t_2|n_1t_1), \tag{I.16}$$

which using (I.9) and replacing in (I.15) gives [2]

$$C(t_2,t_1) = \sum_{n_1}^{\infty} n_1^2 P(n_1t_1|n_0t_0) e^{(\sigma-\kappa)(t_2-t_1)}, \tag{I.17}$$

and since $\sum_{n_1}^{\infty} n_1^2 P(n_1t_1|n_0t_0) = \langle n_1^2\rangle$, using the result for the variance yields

$$C(t_2,t_1) = (\mathrm{var}_n(t_1) + \langle n_1\rangle^2)\, e^{(\sigma-\kappa)(t_2-t_1)}, \tag{I.18}$$

or

$$C(t_2,t_1) = \left[\frac{\kappa+\sigma}{\kappa-\sigma}\, n_0\, e^{(\sigma-\kappa)t_1}\,(1-e^{(\sigma-\kappa)t_1}) + n_0^2\, e^{2(\sigma-\kappa)t_1}\right] e^{(\sigma-\kappa)(t_2-t_1)}. \tag{I.19}$$

For $\sigma = \kappa$, the same procedure gives

$$C(t_2,t_1) = n_0^2 + 2n_0\kappa t_1. \tag{I.20}$$

The correlation function of the *fluctuations* $\delta n(t) = n(t) - \langle n(t)\rangle$, or *connected* time correlation function is

$$C_c(t_2,t_1) = C(t_2,t_1) - \langle n_1\rangle\langle n_2\rangle.$$

This function is zero when the variables are uncorrelated (which is typically expected to happen for $t \to \infty$, although this is not the case in the growth phase, see below). For this reason this function is often more useful than the time correlation function (2.1), which for uncorrelated variables takes the value $\langle n_1\rangle\langle n_2\rangle$. In the case $\sigma \neq \kappa$ the connected time correlation function is

$$\begin{aligned} C_c(t_2,t_1) &= \left[\frac{\kappa+\sigma}{\kappa-\sigma}\, n_0\, e^{(\sigma-\kappa)t_1}\, (1-e^{(\sigma-\kappa)t_1})\right] e^{(\sigma-\kappa)(t_2-t_1)} \\ &= \mathrm{var}_n(t_1)\, e^{(\sigma-\kappa)(t_2-t_1)}, \end{aligned} \tag{I.21}$$

and for $\sigma = \kappa$:

$$C_c(t_2,t_1) = 2n_0\kappa t_1 \tag{I.22}$$

Normalizing $C_c(t_2,t_1)$ by $\sqrt{\mathrm{var}_n(t_1)\mathrm{var}_n(t_2)}$ one obtains the *Pearson correlation:*

$$\rho = \frac{C_c(t_2,t_1)}{\sqrt{\mathrm{var}_n(t_1)\mathrm{var}_n(t_2)}}. \tag{I.23}$$

In our case this gives

$$\rho(t_1,t_2) = \sqrt{\frac{\mathrm{var}_n(t_1)}{\mathrm{var}_n(t_2)}}\, e^{(\sigma-\kappa)(t_2-t_1)}, \qquad \text{if } \sigma \neq \kappa, \tag{I.24}$$

$$\rho(t_1,t_2) = \sqrt{\frac{t_1}{t_2}}, \qquad \text{if } \sigma = \kappa. \tag{I.25}$$

Note the power law behaviour for $\sigma = \kappa$, which corresponds to an infinite correlation time, a feature of critical dynamics. Note also that when $t_2 \to \infty$ we have for $\sigma \leq \kappa$ that

$$\lim_{t_2\to\infty} \rho(t_1,t_2) = 0, \tag{I.26}$$

i.e. the system decorrelates, while if $\sigma > \kappa$

$$\lim_{t_2\to\infty} \rho(t_1,t_2) = \sqrt{1-e^{-(\sigma-\kappa)t_1}}, \tag{I.27}$$

i.e. the limit depends on the initial time, and the system remains forever correlated.

#### II. NON-MARKOVIAN MODEL

To take into account the cell cycle, we formulate a model with the same assumptions as before (equivalent and independent families descending from each individual), but introduce the age, or proper time of each cell, $\tau$, that is the time elapsed since the division that gave rise to it, and a $\tau$-dependent proliferation rate $\gamma(\tau)$ (we will not consider apoptosis in this case). Following van Kampen [4], one can write an integral equation for the generating function,

$$g(x,t) - x = \int_0^t \left[g^2(x,t-t') - x\right] \gamma(t')w(t')\, dt', \tag{II.1}$$

which is valid for the initial condition $n(t=0) = 1$, and where $w(t)$ is the probability that a cell reaches the age $\tau$ without dividing,

$$w(\tau) = e^{-\int_0^\tau \gamma(t')dt'}. \tag{II.2}$$

Using (I.3) and (I.4), we deduce from (II.1) integral equations for the mean and mean square of population,

$$\langle n(t)\rangle - 1 = \int_0^t \left[2\langle n(t')\rangle - 1\right] w(t-t')\gamma(t-t')dt', \tag{II.3a}$$

$$\langle n^2(t)\rangle - \langle n(t)\rangle = \int_0^t 2\left[\langle n(t-t')\rangle^2 + \langle n^2(t-t')\rangle - \langle n(t-t')\rangle\right] \gamma(t')w(t')\, dt'. \tag{II.3b}$$

Below we solve these for the choice

$$\gamma(\tau) = \gamma_0 H(\tau - T), \tag{II.4}$$

where $H(t)$ is the Heaviside function. In other words, we introduce a cell cycle with a refractory period $T$ such that no proliferation is possible for $\tau < T$, and it happens with constant rate $\gamma_0$ for $\tau \geq T$. Using this form of $\gamma(\tau)$ in (II.2) gives

$$w(\tau) = e^{-\gamma_0 H(\tau-T)(\tau-T)}. \tag{II.5}$$

##### A. Population mean

Equation (II.3a) can be solved introducing the Laplace transforms

$$n(s) = \int_0^\infty e^{-st}\langle n(t)\rangle\, dt, \qquad w(s) = \int_0^\infty e^{-st} w(t)\, dt, \qquad y(s) = \int_0^\infty e^{-st} w(t)\gamma(t)\, dt. \tag{II.6}$$

One finds

$$n(s) = \frac{1}{s}\frac{1-y(s)}{1-2y(s)} = \frac{w(s)}{1-2y(s)}, \tag{II.7}$$

where the second equality follows because $dw/dt = -\gamma(t)w(t)$, so that $w(s) = (1-y(s))/s$.

As a check of this result one can set $T = 0$ in (II.4) (i.e. $\gamma(\tau) = \gamma_0$) and recover the Markovian result. Since in this case $y(s) = \gamma_0/(s+\gamma_0)$, it follows that $n(s) = 1/(s-\gamma_0)$ and $\langle n(t)\rangle = e^{\gamma_0 t}$.

When $T > 0$, $w(s)$ and $y(s)$ are found to be

$$w(s) = \frac{1-e^{-Ts}}{s} + \frac{e^{-Ts}}{s+\gamma_0}, \qquad y(s) = \gamma_0 \frac{e^{-Ts}}{s+\gamma_0}, \tag{II.8}$$

so that the Laplace-space solution (II.7) reads

$$n(s) = \frac{1}{s} + \frac{\gamma_0 e^{-Ts}}{s\phi(s)}, \qquad \phi(s) = s + \gamma_0\left(1 - 2e^{-Ts}\right). \tag{II.9}$$

To find an expression for the mean population in the time domain we need to compute the inverse Laplace transform,

$$\langle n(t)\rangle = \frac{1}{2\pi i}\int_{\alpha-i\infty}^{\alpha+i\infty}\left[\frac{1}{s} + \frac{\gamma_0 e^{-Ts}}{s\phi(s)}\right] e^{st}\, ds, \tag{II.10}$$

where $\alpha$ is a real number such that the integration contour lies to the right of all the poles of $n(s)$.

We will evaluate the inverse transform using the residue theorem, so we need to find the poles of $n(s)$. These are located at $s = 0$ and at the zeros of $\phi(s)$. It is easy to check that the zeros must appear in complex conjugate pairs, i.e. if $\phi(\hat{s}) = 0$ then $\phi(\hat{s}^*) = 0$. They can be written in terms of the Lambert, or product log, function $W_k(z)$. For each integer $k$, $W_k(z)$ is a branch of the converse of $f(z) = ze^z$ (with complex $z$), i.e. if $v = W_k(z)$ , then $ve^v = z$. One finds a zero of $\phi(s)$ for each branch of $W_k(z)$,

$$\hat{s}_k = -\gamma_0 + \frac{1}{T}W_k\left(2T\gamma_0 e^{T\gamma_0}\right). \tag{II.11}$$

Since the argument of $W_k$ is real, there is one real zero, $\hat{s}_0$, and $s_{-k} = s_k^*$. Furthermore it holds that if $|j| > |k|$, then $\mathrm{Re}\, s_j < \mathrm{Re}\, s_k$.

Since $\phi(0) = -\gamma_0 \neq 0$, the integrand of (II.10) has simple poles at $s = 0$, $s = s_k$, and the contour must run through a line parallel to the imaginary axes, $\alpha \geq \max(0, \hat{s}_0)$. Using the residue theorem we have finally

$$\langle n(t)\rangle = 1 + H(t-T)\left[-1 + \sum_{k=-\infty}^{\infty}\frac{\gamma_0}{\hat{s}_k}\frac{e^{\hat{s}_k(t-T)}}{1+T(\gamma_0+\hat{s}_k)}\right]. \tag{II.12}$$

for the initial condition $n(t=0) = 1$. If instead the system starts with $n_0$ *synchronised* cells at $t = 0$, then we have simply

$$\langle n(t)\rangle = n_0 + n_0 H(t-T)\left[-1 + \sum_{k=-\infty}^{\infty}\frac{\gamma_0}{\hat{s}_k}\frac{e^{\hat{s}_k(t-T)}}{1+T(\gamma_0+\hat{s}_k)}\right]. \tag{II.13}$$

##### B. Variance

Introducing the Laplace transforms $\langle n(t)\rangle^2$ and $\langle n^2(t)\rangle$, respectively $m(s)$ and $n_2(s)$, equation (II.3b) becomes, in the Laplace domain,

$$n_2(s) - n(s) = 2\left[m(s) + n_2(s) - n(s)\right] y(s), \tag{II.14}$$

and solving for $n_2(s)$ one can write the variance in the Laplace domain,

$$\text{var}_n(s) = n(s) - m(s) + m(s)\frac{2y(s)}{1-2y(s)} = n(s) - m(s) + 2\gamma_0 e^{-sT}\frac{m(s)}{\phi(s)}. \tag{II.15}$$

To transform to the time domain it is convenient to define $\psi(s) = \frac{1}{\phi(s)}$ , $h(s) = \psi(s)m(s)$, so that

$$\text{var}_n(t) = \langle n(t)\rangle - \langle n(t)\rangle^2 + 2\gamma_0 H(t-T)h(t-T), \tag{II.16}$$

where in the time domain $h(t)$ can be written as a convolution:

$$h(t) = \int_0^t \psi(t-t')\langle n(t')\rangle^2 \, dt'. \tag{II.17}$$

This way of expressing the solution is convenient because we already have $\langle n(t)\rangle$ in the time domain (II.13), and $\psi(t)$ can be obtained by the residue theorem using the same zeros $s_k$ (it has the same poles as $n(s)$ except for $s = 0$), giving

$$h(t) = \int_0^t \sum_k \frac{e^{s_k(t-t')}}{\phi'(s_k)}\left\{H(t-T) + H(t-T)\gamma_0^2 \sum_{l,k} \frac{e^{(s_k+s_l)t'}}{s_k s_l \phi'(s_k)\phi'(s_l)}\right\} dt' \tag{II.18}$$

with $\phi'(s)$ the derivative of $\phi(s)$. Performing the integral and collecting everything into (II.16), we can write the final expression for the variance:

$$\begin{aligned}\text{var}_n(t) = n_0^2 H(t-T)&\left\{3\gamma_0 \sum_k \frac{e^{s_k(t-T)}}{s_k\phi'(s_k)} - \gamma_0^2 \sum_{k,l} \frac{e^{(s_k+s_l)(t-T)}}{s_k s_l \phi'(s_k)\phi'(s_l)}\right\} + \\ 2\gamma_0 n_0^2 H(t-2T)&\left\{\gamma_0^2 \sum_{k,l,m} \frac{e^{(s_l+s_m)(t-T)} - e^{(s_l+s_m)T} - e^{s_k(t-2T)}}{(s_l+s_m-s_k)s_m s_l \phi'(s_m)\phi'(s_l)\phi'(s_k)} - \sum_k \frac{e^{s_k(t-T)}}{s_k\phi'(s_k)}\right\}.\end{aligned} \tag{II.19}$$

##### C. Numerical evaluation of mean and variance

To numerically evaluate the mean we used the Eq. (II.13) directly, including $\sim 10$ poles in the sum (always taking the complex conjugate pairs so as to obtain a real result). We used a script in Julia [5], computing the poles $\hat{s}_k$, Eq. (II.11) using Julia's `lambertw` function from module `LambertW` [6]. For the evaluation of the variance we used Eq. (II.16), performing the convolution in (II.17) numerically with Julia's `conv` function from the DSP module [7]. The explicit expression (II.19) for the variance is tricky to evaluate numerically due to cancellations in the triple sum of the second term; an accurate numerical evaluation using this formula requires extended numerical precision.

#### III. NUMERICAL SIMULATIONS

For the numerical simulation we defined a discrete-time version of the contact processes described above. The time step is $\Delta t$ and at each step each cell can undergo division with probability $P_\sigma$ or apoptosis with probability $P_\kappa$, with $P_\sigma + P_\kappa \leq 1$. The refractory period $T$ is a multiple of $\Delta t$, and record is kept of whether each cell's age is larger than $T$. At each step and for each cell a random number $p$ uniform in $[0,1]$ is drawn. If $p \leq P_\kappa$, the cell is removed, while if $P_\kappa < p \leq P_\kappa + P_\sigma$ and the cell's age is larger than $T$, the cell divides. The simulations to compare with the Markovian contact process were done with $T = 0$, while for the comparison with the non-Markovian process we set $P_\kappa = 0$. To perform quantitative comparisons, the probabilities are set in terms of the transition rates as

$$P_\sigma = 1 - e^{-\sigma\Delta t}, \qquad P_\kappa = 1 - e^{-\kappa\Delta t}. \tag{III.1}$$

For the non-Markovian case, $\gamma_0$ is used in place of $\sigma$. The model was implemented in Python [8].

#### IV. SUPPLEMENTARY FIGURES

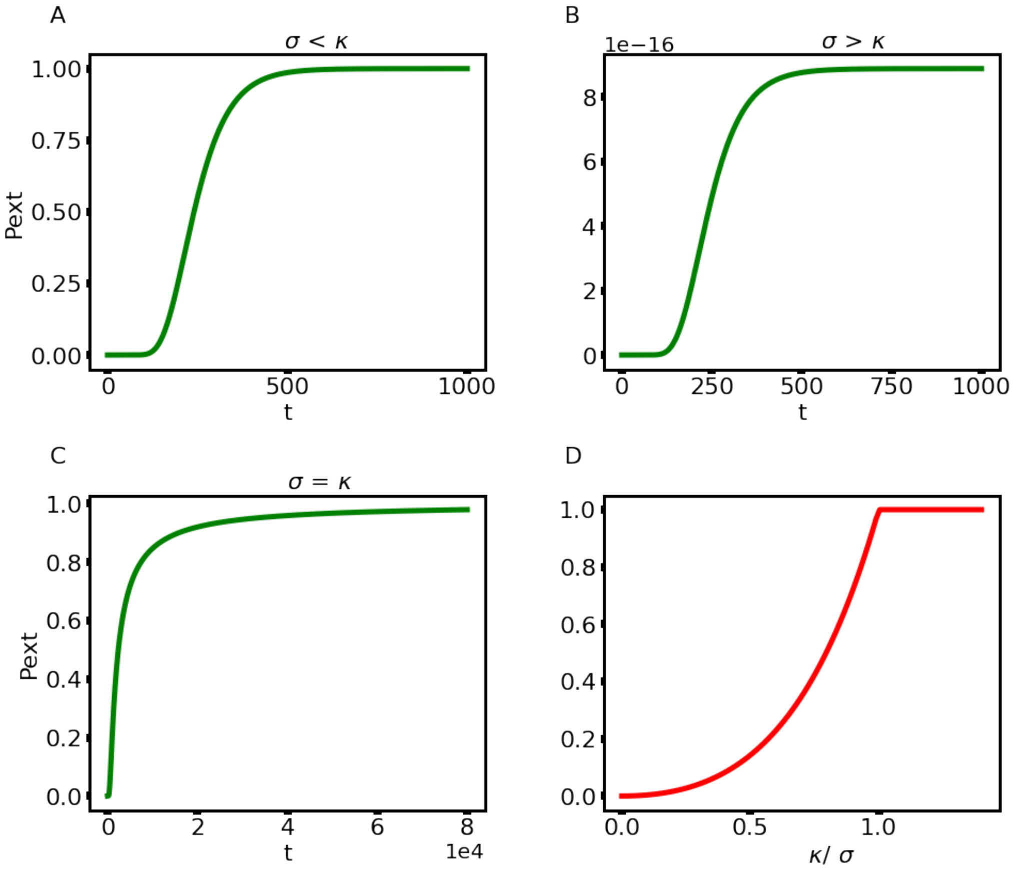


Supplementary figure 1. **Extinction probability for the different regimes of the Markovian model.** The extinction probability $P_e(t)$ (the probability that the system becomes extinct at or before $t$) starting with $n_0$ =50 for the decline phase (A), growth phase (B) and homeostatic state (C). When $t \to \infty$, $P_e$ tends to 1 in the homeostatic and decline regimes, and to a value $(\kappa/\sigma)^{n_0} < 1$ in the growth regime (D). Note the singularity at $\kappa/\sigma = 1$.

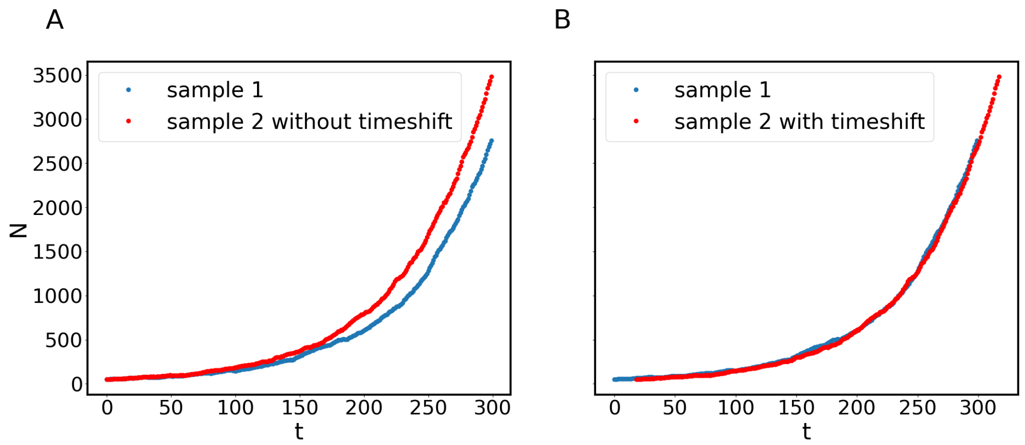


Supplementary figure 2. **The growth of two different samples is approximately related by a time shift.** A) Number of cells vs. time for two samples of the numerical simulation of the Markovian model. B) Same curves with the curve of sample 2 shifted in time by 18 time steps. This shows that the exponential growth in both samples is essentially amplifying the initial small fluctuations among samples, so that growth in different samples is approximately related by a time shift. After the first few steps, if one sample has a larger population than another one, it is extremely unlikely that the size relation will invert later in time.

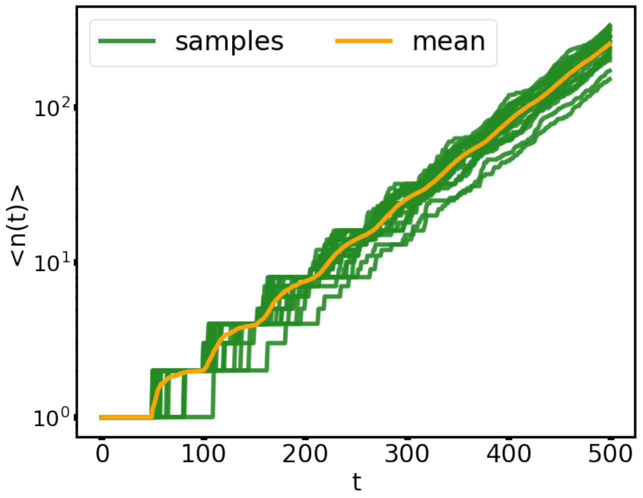


Supplementary figure 3. **The temporal evolution of the number of cells shows damped oscillatory behaviour superimposed over the growth.** In orange, temporal evolution (500 simulation steps) of the average number of cells over 200 samples of a tissue. In green temporal evolution of 20 samples.


[1] U. C. Täuber, *Critical Dynamics* (Cambridge University Press, 2014) pp. 70–74.

[2] I. Pázsit and L. Pál, *Neutron Fluctuations: A Treatise on the Theory of Branching Processes* (Elsevier, Amsterdam, 2008).

[3] R. Garcia-Millan, J. Pausch, B. Walter, and G. Pruessner, Field-theoretic approach to the universality of branching processes, Phys. Rev. E **98**, 062107 (2018), arxiv:1808.08418.

[4] N. V. Kampen, *Stochastic Processes in Physics and Chemistry*, 3rd ed. (North Holland, 2007).

[5] J. Bezanson, A. Edelman, S. Karpinski, and V. B. Shah, Julia: A fresh approach to numerical computing, SIAM review **59**, 65 (2017).

[6] Docstrings — lambertw.jl, `https://docs.juliahub.com/LambertW/7mpiq/0.4.5/autodocs/`, accessed: 2023-04-20.

[7] Docstrings — dsp.jl, `https://docs.juliadsp.org/stable/convolutions/`, accessed: 2023-04-20.

[8] G. Van Rossum and F. L. Drake, *Python 3 Reference Manual* (CreateSpace, Scotts Valley, CA, 2009).
